# Assessing the efficacy of antibiotic treatment for the creation of axenic earthworms

**DOI:** 10.1101/2021.04.16.440126

**Authors:** Henny O. Omosigho, Elmer Swart, Tom P. Sizmur, Dave J. Spurgeon, Claus Svendsen, Liz J. Shaw

## Abstract

Earthworms are an integral part of soil ecosystems, especially for their role in soil functions such as organic matter (OM) decomposition and nutrient cycling. Earthworms and microorganisms are interdependent, and a considerable portion of the contribution earthworms make to influencing OM fate is through interactions with microorganisms. However, the importance of the earthworm-associated microbiome is not fully understood, because it is difficult to separate the direct influence of the earthworms from the indirect influence of their microbiome. Here, we evaluated an antibiotic-based procedure for producing axenic ecologically-contrasting earthworm species (*E. fetida, L. terrestris, A. chlorotica*) as the first step towards soil studies aimed at understanding the importance of the earthworm microbiome for host health and function. Individual earthworms were exposed to antibiotics: cycloheximide (150 μg ml^−1^), ampicillin (100 μg ml^−1^), ciprofloxacin (50 μg ml^−1^), nalidixic acid (50 μg ml^−1^), and gentamicin (50 μg ml^−1^) either singly or in a cocktail via culture (96 h) in a semi-solid agar carrier. Compared to the non-antibiotic treated control, the cocktail (for all three species) and ciprofloxacin (for *E. fetida* and *A. chlorotica*) treatments significantly reduced (P<0.05) culturable microbial abundance on nutrient agar and potato dextrose agar. The microbial counts were reduced to below detection (<50 CFU individual^−1^) for *E. fetida* and *A. chlorotica* receiving the cocktail. Illumina 16S rDNA amplicon sequence analysis of culturable *L. terrestris* -associated bacteria showed that antibiotic treatment influenced community composition revealing putative sensitive (*Comomonas, Kosakonia* and *Sphingobacterium*) and insensitive (*Aeromonas, Pseudochrobactrum*) taxa. Overall, we report a rapid, with minimal earthworm-handling, process of creating ‘axenic’ *E. fetida* and *A. chlorotica* individuals or *L. terrestris* with a suppressed microbiome as a tool to be used in future ecological studies of earthworm microbial interactions affecting host health and function.

## 1. Introduction

Earthworms are one of the most dominant soil invertebrates in terms of biomass [1,2] and are frequently referred to as ‘ecosystem engineers’ due to their effects on soil structure and nutrient availability [3]. Earthworms have been classified into three main ecological categories (epigeic, endogeic and anecic groups) by Bouché (1977) [4] based on ecological and morphological characteristics as well as their vertical distribution in the soil profile [4–6]. Epigeic species are surface dwelling, non-burrowing and consume decaying plant residues on the soil surface. Anecic worms build permanent vertical burrows but feed on plant litters at the surface or dragged into burrows to be pre-decomposed by microorganisms; endogeic worms inhabit and feed in organo-mineral and deeper mineral horizons [2,4]. Recently, Bottinelli et al. 2020 [6] applied a numerical approach to the classification of earthworms to the ecological categories. This approach enabled a given species to be defined by three percentages of membership to the three main categories and allowed for species to belong to supplemental intermediary categories (e.g., epi-anecic or epi-endo-anecic).

Earthworms are major players in determining soil organic matter (SOM) dynamics [7,8]. Earthworms not only stimulate organic matter (OM) decomposition, but they also promote SOM stabilization within soil aggregates [9,10]. Decomposition is enhanced both by increasing the access of microbial decomposers to OM substrates through mixing and fragmentation of litter [9,11–14] and by stimulating the activity of the ingested soil-derived earthworm gut microbiome, which accelerates the breakdown of earthworm-ingested OM during gut passage. This latter is referred to as ‘the sleeping beauty paradox’ [3,15]. It involves the production of intestinal C-rich mucus (‘the kiss’) by the earthworm (‘Prince Charming’). This process awakens ingested dormant microflora (‘sleeping beauties’) and thereby increases the decomposition of ingested organic matter because of a ‘priming’ effect [15–18].

It has long been suggested that most earthworm species are not capable of secreting the full set of enzymes that are required for the depolymerization of plant-derived polymers. Whilst the possession of endogenous endocellulase genes by some earthworm species has been reported [19], indicating the ability to digest cellulose, it is thought that even when earthworms can produce endocellulase, their ability to digest and acquire nutrients from plant litter lies fundamentally in their relationship with microorganisms [20]. This is because efficient degradation of a complex polymer such as lignocellulose requires the synergistic action of suites of enzymes, such as hemicellulase, endocellulase, lignin peroxidase and exocellulase, that are primarily secreted by microorganisms [21]. The role of the aforementioned ‘kiss’ may therefore be to stimulate microbial depolymerase production during gut passage to aid acquisition of nutrients from ingested plant litter. However, depolymerase activity in soil is a function of recently secreted enzymes, and those produced in the past and stabilized through association with the soil matrix [22,23]. Therefore, it is not clear if earthworms rely on the microbial production of enzymes during gut transit, or, if already produced enzymes (before ingestion) are sufficient for complete depolymerisation. In the latter case, earthworms would not depend on ingested microorganisms themselves, but only on their pre-produced enzymes that were obtained through ingestion.

In addition to a role of an active, soil-derived, gut microbiome for host nutrition, it is possible that the earthworm microbiome is also vital for other purposes. For example, many studies have suggested that gut microbiomes of various hosts such as humans, *Drosophila melanogaster* (fruit fly), *Riptortus pedetris* (bean bug) and termites, play essential roles in different physiological processes. This includes immunity [24–27], reproduction [28], and resistance to pesticide-induced stress [29]. The earthworm gut microbiome, and indeed the microbiome associated with the other organs (such as skin and the nephridia), may confer additional functions that extend beyond roles in digestion and provision of nutrients to the host such as functions that affect host sexual maturity and reproduction [30,31].

Despite the uncertainties regarding the role of the earthworm microbiome in providing nutritional and non-nutritional benefits to the host, comprehensive studies on this topic, and on the role played by the earthworm host-microbiome interaction for ecosystem processes, are lacking. These uncertainties are due to our inability to separate the contribution of ingested and ‘native’ microbes to host processes. Therefore, we need a method to eliminate the role played by the ‘native’ microbiome to allow the understanding of the contributions of the host, the microbiome (and their interactions) to functional effects.

Previous studies have attempted to produce axenic earthworm cultures through the passage of individual animals via sterile solutions or suspensions containing antibiotics, both single antibiotics and cocktail of antibiotics [32,33]. These studies used *E. fetida* as the ‘model’ organism; presumably because it can easily be reared on a variety of organic substrates [34] using standard protocols [35]. However, *E. fetida* is not a typical soil dwelling earthworm species [35], preferring organic-rich habitats. Hence to understand microbiome effects, there is a need to extend studies to other species of earthworm occupying different niches within the soil.

In this present study, we developed and evaluated an antibiotic-based procedure for producing axenic specimens of earthworms belonging to the epi- anecic (*L. terrestris*) and epi-endo-anecic (*A. chlorotica*) ecological groupings as well as *E. fetida* as a comparatively well-studied comparison. The study, thus, provides a first step towards understanding the importance of the earthworm non-transient microbiome for earthworm health and ecological functional roles. We evaluated the effects of antifungal and anti-bacterial antibiotic treatments (individually and in a cocktail) on culturable earthworm-associated microbial numbers. To further understand how antibiotic exposure influenced the *L. terrestris*-associated culturable bacterial diversity, we used 16S rDNA amplicon sequencing. This provided insights into the taxa specific changes associated with specific treatment knockdowns.

## 2. Material and methods

### 2.1 Earthworm collection and culture

*E. fetida* and *L. terrestris* were purchased from Worms Direct (Essex, UK). *A. chlorotica* specimens were collected from the University of Reading dairy farm at Shinfield (grid reference 51.408580, −0.927353) by hand sorting for adult *A. chlorotica*, identified using the guide of Sherlock [36]. Identified earthworms were washed with autoclaved de-ionised water before transport back to the laboratory in a cool box. Each earthworm species was acclimated to laboratory conditions in the dark at 20 ± 2 °C for two weeks [37,38] prior to the start of the experiment in a culture made from Kettering loam and Irish moss peat (2:1 ratio) [39] and the earthworms were fed Irish moss peat at approximately 1 g earthworm^−1^ week^−1^ after one week of acclimation [38].

### 2.2 Antibiotic exposure

The adult earthworm individuals selected for antibiotic exposure were responsive to stimuli and visibly healthy. Selected individuals were of similar sizes (within the same species) to avoid the potential for size-specific effects. Following initial depuration (48 h on moist filter paper in the dark), single earthworm specimens were incubated in Duran bottles of either 250 ml (*E. fetida* and *A. chlorotica*) or 500 ml (*L. terrestris*) in volume, containing either 50 ml (*E. fetida* and *A. chlorotica*) or 150 ml (*L. terrestris*) of sterile 0.65 % (m/v) technical agar medium (Fisher Scientific, UK). The technical agar concentration used resulted in a medium that, as determined in a preliminary experiment, was of a consistency that allowed the earthworms to burrow within the agar. The agar volume ensured that there was an agar depth of at least 10 cm, as this was found to be a suitable depth, especially for the anecic earthworms, to form vertical burrows [40]. The agar medium was supplemented with antibiotics (Sigma-Aldrich) added individually or as a cocktail of the five antibiotics in the concentrations shown in Table 1. The concentration of each antibiotic in the cocktail was the same as the concentration used when a single antibiotic was applied. Hence when combined this treatment provides both a more complex and greater total antibiotic exposure treatment. The antibiotics were chosen to represent different classes based on mechanism of action [41]. Antibiotics that were not purchased as already-made solutions but in solid form were dissolved in either 0.1 M hydrochloric acid (ciprofloxacin) or distilled water (nalidixic acid) as required to make up stock solutions.

**Table 1.** Antibiotic types and concentrations used to amend agar media for the production of ‘axenic’ earthworms

| Antibiotic | Antibiotic concentration ($\mu\text{g ml}^{-1}$ agar) |
| --- | --- |
| Cycloheximide | 150<sup>a</sup> |
| Ampicillin | 100 |
| Ciprofloxacin | 50 |
| Nalidixic acid | 50<sup>a</sup> |
| Gentamicin | 50<sup>a</sup> |

<sup>a</sup> Antibiotic concentration used in Whiston and Seal (1988)

For each earthworm species, triplicate samples for each antibiotic treatment were incubated at 20 ± 2 °C in darkness for 96 hours following earthworm addition. Control samples with technical agar but without antibiotics added were included (n = 3). The bottles were covered with aluminium foil to prevent earthworm escape, with pin holes in the cover to ensure aeration.

### 2.3 Assessment of the abundance and diversity of earthworm-associated culturable microorganisms

#### 2.3.1 Microbial abundance

After 96-hours of antibiotic exposure, the earthworms were removed from the agar medium with sterile tweezers. No earthworm mortality was recorded during the incubation period and all earthworms had burrowed and were responsive to a physical stimulus. Following removal from the antibiotic medium, earthworms were washed with autoclaved de-ionised water and placed in 10 ml sterile centrifuge tubes. Earthworms were euthanised when placed in a 4°C cold room for 1 hr and then crushed using sterile glass rods. One ml of autoclaved de-ionised water was added to the tube, followed by vigorous shaking (250-rev min^−1^ for 2 min). The resulting suspension was serially diluted in triplicate with autoclaved de-ionised water in a ten-fold dilution series (10^0^, 10^−1^, 10^−2^, 10^−3^, 10^−4^, 10^−5^, 10^−6^, 10^−7^ and 10^−8^). To determine the number of colony-forming units (CFUs) of culturable earthworm-associated microorganisms, 20 μl (*E. fetida* and *A. chlorotica*) or 200 μl (*L. terrestris*) of the dilutions were plated on to agar plates following [42] or using the spread plate method, respectively. Nutrient agar (NA), that predominantly favours bacterial growth, and potato dextrose agar (PDA), normally used for culturing fungi, were the agar media used. The agar plates were incubated in the dark at 26 °C under oxic conditions. The emerging colonies were observed after 24 hrs and then at two weeks when the colonies were counted. The limit of detection of the plate count method was determined using the volume plated and the dilution factor [43].

#### 2.3.2 DNA Extraction, 16S-rDNA sequencing

Out of the three earthworm species studied, *L. terrestris* (as the only species that had CFUs above detection limits for all antibiotic treatments and both agars) was carried forward for DNA-based analysis of associated microorganisms that were cultured on plates arising from the dilution spread plate estimation of microbial abundance.

For each antibiotic treatment, earthworm individual and agar type, colonies growing across all dilutions were harvested using a sterilised cell scraper. Harvested cells from each plate were initially suspended in 1 ml sterile de-ionised water in a 2 ml centrifuge tube and then the different dilutions of the same replicates were pooled and stored at −20 °C prior to DNA extraction.

Total genomic DNA was extracted from the samples using DNeasy Ultraclean Microbial Kit (Qiagen) according to the manufacturer’s protocol. The quality and concentration of the extracted DNA sample was measured using a NanoDrop spectrophotometer (ND-2000/2000c, NanoDrop Technologies).

A ~550 bp fragment of the V3-V4 hypervariable region of the bacterial 16S-rRNA gene was amplified by PCR with 5’-CCTACGGGAGGCAGCAG-3’ as the forward primer and 5’-GGACTACHVGGGTWTCTAAT-3’ as the reverse primer. Each reaction was done in a 50 μl reaction using four ng of genomic DNA. Each sample was dual index barcoded following [44]. The amplification thermal cycling consisted of an initial denaturing step at 95 °C for 2 minutes, followed by 30 cycles of denaturation at 95 °C for 30 seconds, annealing at 55 °C for 15 seconds and extension at 72 °C for 40 seconds, with a final extension step at 72 °C for 10 minutes. All PCR reactions were performed using Q5^®^ High-Fidelity DNA Polymerase (New England BioLabs, USA). Quality and verification of fragment size was performed using gel electrophoresis. Samples were normalised using a SequalPrep Normalisation Plate Kit (Thermo Fisher Scientific, UK) and subsequently pooled. The pooled samples were subsequently run on a 1.2% agarose gel and a ~550 bp fragment was gel extracted using a QIAquick Gel Extraction Kit (Qiagen, the Netherlands). The gel extracted samples were quantified using a Qubit HS DNA Kit (Thermo Fisher Scientific, USA) and diluted to 7 pM using HT buffer. The final library was the run with 10% PhiX using a MiSeq Reagent Kit v3 (600 cycles) on a MiSeq (Illumina, USA). Nucleotide sequence data have been submitted to NCBI and are available under submission number SUB9306713 as part of bioproject ID PRJNA715719.

### 2.4 Statistical and bioinformatics analyses

The effect of antibiotic treatment on the number of CFUs for each earthworm species (*E. fetida, A. chlorotica*, and *L. terrestris* was assessed using one-way analysis of variance (ANOVA) followed by post hoc Tukey comparisons, where appropriate (P<0.05). Normality of residuals (Anderson-Darling test) and equal variance (Levine’s test) assumptions were verified, and data was square root transformed where required. All analyses for the plate count data were performed with Minitab 19.1.1.

MiSeq reads were demultiplexed using BaseSpace (Illumina, USA). Amplicon sequence variant (ASV) tables were generated using the DADA2 pipeline [45]. Briefly, in this procedure, the forward and the reverse reads were inspected for quality. The quality of the reverse reads was below 30 from 200 bases onwards. This prevented sufficient merging of the forward and reversed reads, and hence only the forward reads were used in further analysis. The last ten bases of the forward reads were trimmed, and trimmed reads were subsequently filtered applying a maxN, maxEE and truncQ value of 0, 2 and 2, respectively. After sample inference, reads were subjected to chimera removal. Filtering of low-quality reads and removal of chimera lead to removal of on average 18% of the forward reads per sample. Taxonomy was assigned using the Silva version 132 dataset [46].

ASVs assigned to eukaryotes, archaea, chloroplasts, and mitochondria or to an unknown phylum or kingdom were removed from the dataset.

All statistical analyses of ASVs data and data visualisations were performed in R v.3.6.3 [47]. The diversity analysis was carried out using the packages ‘phyloseq’ (McMurdie and Holmes, 2013) and ‘vegan’ [48]. Observed and Chao1 richness and phylogenetic diversity measures were used to estimate the alpha diversity. The normality of the dataset was checked using Shapiro-Wilk normality test and the significance of differences between alpha diversity and relative abundance of taxa was evaluated using analysis of variance (ANOVA). For the beta diversity, the principal coordinate analysis (PCoA) based on Jaccard distances that focuses on unique features, regardless of abundance was used to visualise the similarity of individual replicates based on the presence and absence of ASVs. The effect of antibiotic treatment on bacterial community patterns were further analysed by permutation analysis of variance (PERMANOVA) based on Jaccard distances with the Adonis function (999 permutations) of the ‘vegan’ package. The effect of antibiotic treatment on bacterial community patterns was also examined using ANOSIM. ‘VennDiagram’ was used to construct a logical visualisation of relationships between the bacterial genera present in the antibiotic treatments [49].

## 3. Results

### 3.1 Effect of antibiotic treatment on earthworm-associated culturable microbial abundance

The aim was to develop and evaluate an antibiotic-based procedure to eradicate earthworm-associated microorganisms and create axenic earthworms, as far as could be verified using culture-based methods. For the NA plates (Figure 1a, c, e), ANOVA revealed an overall significant effect of antibiotic treatments on the number of colonies forming for *E. fetida* (*P* < 0.001), *A. chlorotica* (*P* < 0.001) and *L. terrestris* (*P* < 0.001). The post hoc Tukey test showed that, when comparing the effect of the individual antibiotic treatments on earthworm-associated microbial abundance across all three earthworm species, cycloheximide and ampicillin had no significant effect on colony formation compared to the non-antibiotic-amended control. All other antibiotic treatments, however, did significantly reduce the microbial burden for at least one earthworm species. The cocktail treatment was the most effective with CFUs on NA reduced to below the limit of detection (<50 CFU/worm) for *E. fetida* and *A. chlorotica* and by more than 2 orders of magnitude for *L. terrestris*. Although the cocktail of antibiotics resulted in the lowest number CFUs, it did not result in a statistically significant different number of CFUs when compared to the ciprofloxacin-only treatment in *E. fetida* and *A. chlorotica* (PDA), although this difference between the cocktail and ciprofloxacin-only treatment was significant for *L. terrestris*.

**Figure 1.**
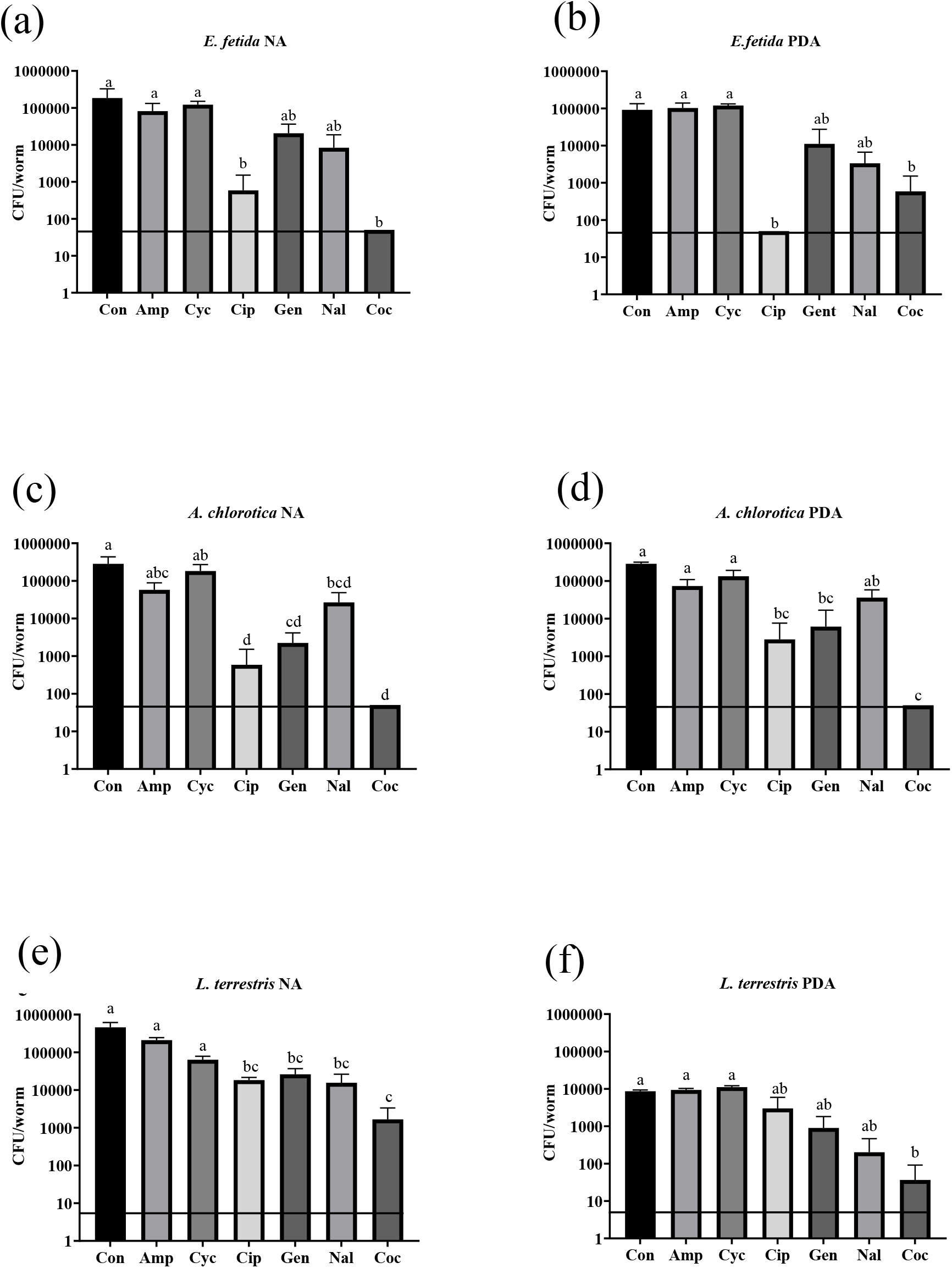
The effect of antibiotic treatment on culturable microbial abundance (Colony Forming Units; CFU) associated with *E. fetida, A. chlorotica* and *L. terrestris* on nutrient agar (NA) and potato dextrose agar (PDA) plates. Con = control; Amp = ampicillin; Cyc = cycloheximide; Cip = ciprofloxacin; Gen = gentamicin; Nal = nalidixic acid; Coc = Cocktail. Values are means ± SE (n = 3). Different letters indicate significant differences between antibiotic treatments for the same agar media and the same earthworm species (Tukey HSD test; α = 0.05). The horizontal line represents the limit of detection for the method of 50 CFU/worm (*E. fetida* and *A. chlorotica*) or 5 CFU/worm (*L. terrestris*).

For the PDA plates (Figure 1 b, d, f), ANOVA indicated a significant effect of antibiotic treatment on the number of CFUs for *E. fetida* (*P* < 0.001), *A. chlorotica* (*P* < 0.001), and *L. terrestris* (*P* < 0.011). Post hoc Tukey test indicated that it was only the cocktail treatment that reduced CFUs compared to control consistently across species. CFU numbers for the cocktail were, however, not statistically different when compared to ciprofloxacin, gentamycin, and (for *E. fetida* and *L. terrestris*) nalidixic acid treatments.

### 3.2 Effect of antibiotic treatment on L. terrestris-associated culturable microbial diversity

#### 3.2.1 16S rDNA amplicon sequencing

Illumina 16S rDNA amplicon sequencing of DNA extracted from colony forming units harvested from NA and PDA dilution series plates from *L. terrestris* generated 1044826 high quality forward reads with an average of 24965 reads per sample. In total 524 ASVs were identified with an average of 31.5 ASVs per sample. Taxonomy was assigned using Silva database version 132 which resulted in the detected bacteria being classed into 10 phyla, 17 classes, 45 orders, 71 families and 143 genera.

#### 3.2.2 Alpha diversity

The observed and estimated (Chao1) ASV richness between individual earthworm replicates had a large variation for control (e.g., for Chao1, the coefficient of variation was 82.31 % for NA plates and 39.5% for PDA plates) and some antibiotic-amended treatments (Figure 2a-d). Against this variable background, one way ANOVA revealed that these alpha diversity measures were not significantly influenced by antibiotic treatment (P>0.05; Figure 2a-d). Similarly, there was no overall effect of antibiotic treatment on Faith’s phylogenetic diversity (P>0.05; Figure 2e, f).

**Figure 2.**
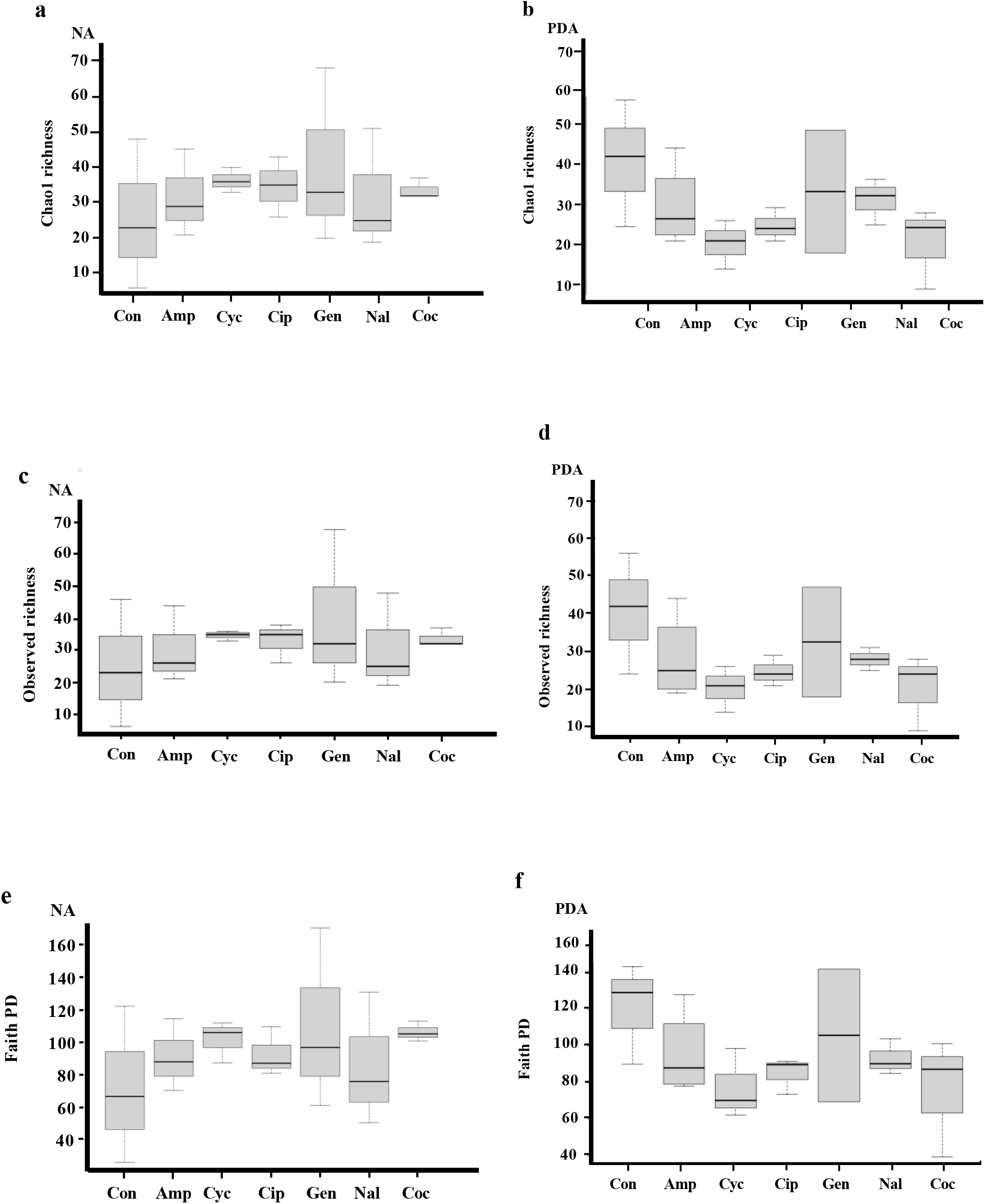
Box plots of ChaoI estimated (a, b) and observed (c, d) Amplicon Sequence Variant (ASV) richness and Faith’s Phylogenetic Diversity (e, f) for *L. terrestris-associated* culturable bacterial communities for control and antibiotic-treated earthworm individuals (n=3) as cultured on nutrient agar (NA; a, c, e) and potato dextrose agar (PDA; b, d, f). Con = control; Amp = ampicillin; Cyc = cycloheximide; Cip = ciprofloxacin; Gen = gentamicin; Nal = nalidixic acid; Coc = Cocktail.

#### 3.2.3 Beta diversity

PCoA based on Jaccard distances was used to visualise the similarity in the data from bacterial community composition for earthworm samples subjected to the different antibiotic treatments (Figure 3). For bacterial communities culturable on NA, the non-antibiotic-treated control samples overlapped with those in the ampicillin- and cycloheximide-treated samples. These clusters appeared distinct from other antibiotic treatment clusters (Figure 3; NA). The PERMANOVA analysis (P = 0.037; [Adonis]) and weakly, the ANOSIM analysis (P = 0.053) supported that the NA-culturable earthworm microbial communities were significantly affected by the antibiotic treatments. The data from the PDA-cultured bacterial community (Figure 3: PDA), also showed that communities from control, ampicillin- and cycloheximide-treated earthworms clustered together and were separated from the clusters of bacterial communities from earthworms treated with nalidixic acid, ciprofloxacin, gentamicin, and cocktail. Both PERMANOVA (*P* = 0.024) and ANOSIM analysis (*P* = 0.009) revealed an overall significant difference between treatment groups. However, for both NA and PDA it is notable that, with few exceptions (ampicillin and control for PDA), individual within-treatment replicates did not group closely together within the ordination space.

**Figure 3.**
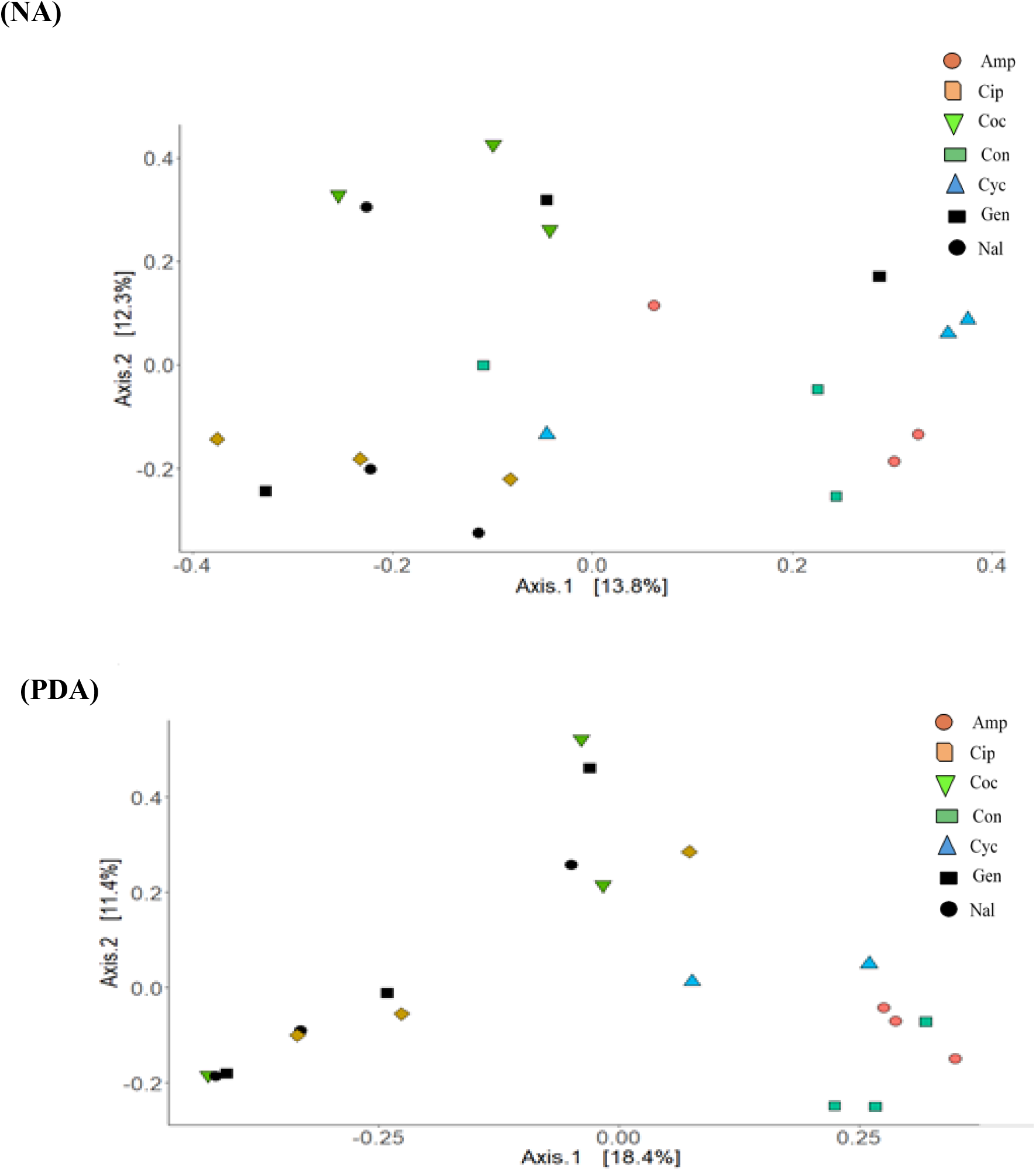
Principal coordinates analysis ordination based on Jaccard distances examining the similarity of composition of culturable bacterial communities for control and antibiotic-treated earthworm individuals (n=3) as determined by 16S rDNA amplicon sequencing of colonies cultured on NA and PDA. Con = control; Amp = ampicillin; Cyc = cycloheximide; Cip = ciprofloxacin; Gen = gentamicin; Nal = nalidixic acid; Coc = Cocktail.

#### 3.2.4 Cultivable shared and unique genera of *L. terrestris* individuals

Given the variability in both alpha and beta diversity at the individual earthworm level (Figure 2; Figure 3), Venn diagrams were used to visualise genera that were unique or common to more than one earthworm individual within the same treatment, with a focus on the non-antibiotic-treated control [to understand the initial variability in the culturable earthworm microbiome (Figure 4a, b)] and the cocktail-treated (Figure 4c, d) earthworm individuals [as the treatment that most significantly impacted culturable *L. terrestris*-associated microbial abundance (Figure 1e, f)]. For nutrient agar, one genera (*Lelliottia*), was consistently detected across control earthworm individuals (Figure 4a). Whilst *Lelliottia* could still be detected in 2 out of 3 cocktail-treated individuals (Figure 4c), two other genera, *Aeromonas* and *Pseudochrobactrum*, were core in cocktail-treated individuals (Figure 4c). Whilst *Aeromonas* was present in the microbiome of two of the NA controls (Figure 4a) (and in all individuals for both control and cocktail treatments for PDA, Figure 4b, d), *Pseudochrobactrum* was not present in any other situation. In addition to *Aeromonas*, 7 other genera were core to control earthworm individuals on PDA (Figure 4b). Out of these, *Pseudomonas, Raoultella* and *Verminephrobacter* were still detected in two of the individuals treated with the antibiotic cocktail (Figure 4d). However, *Comomonas, Kosakonia, Pedobacter* and *Sphingobacterium* could not be detected in cocktail PDA plates (Figure 4d), and, except for *Pedobacter*, were similarly not present in the cocktail treatment for NA plates when they were detected in at least one NA control individual (Figure 4a, c).

**Figure 4.**
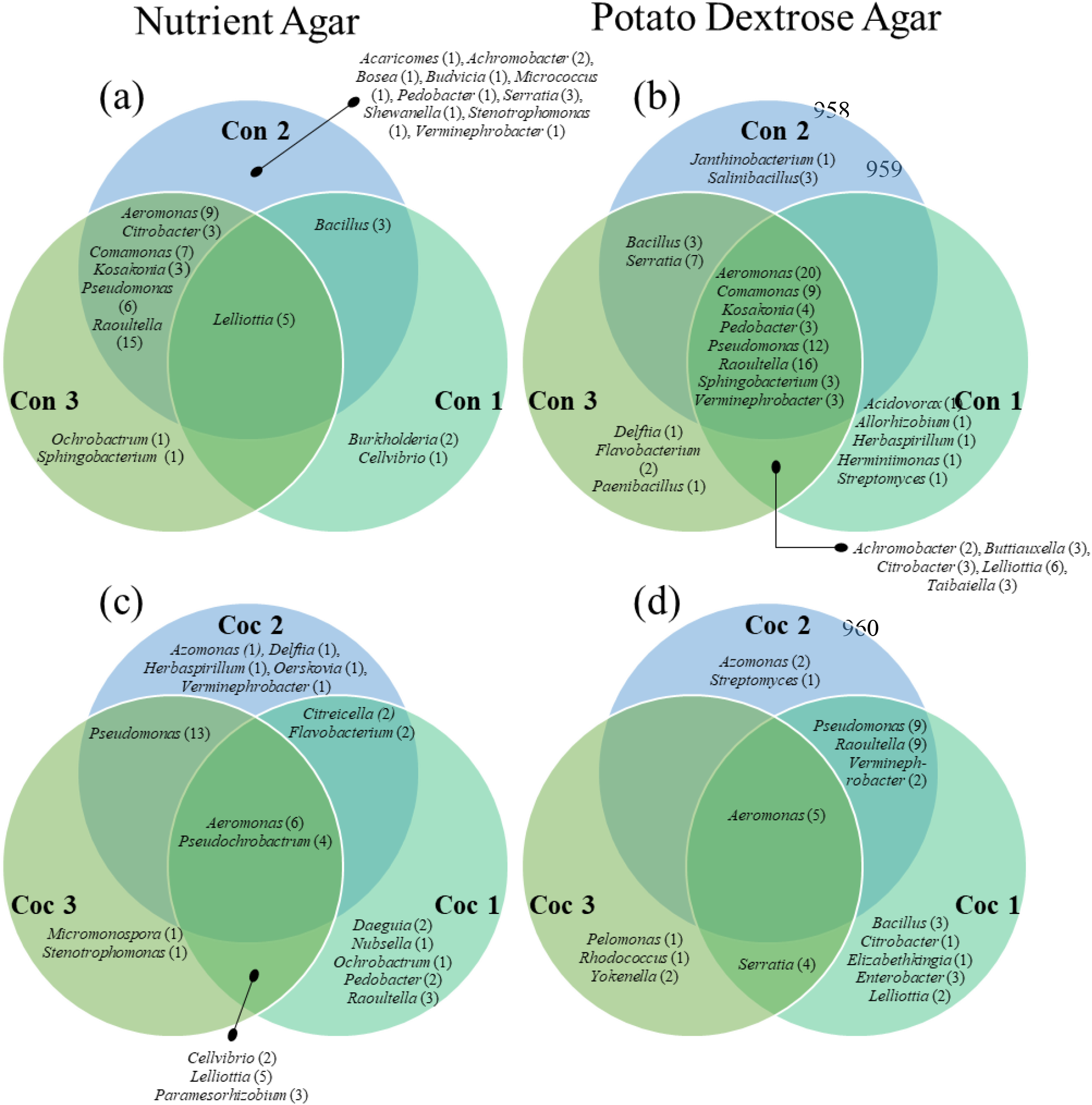
Venn diagram visualisation of genera that are unique or common to more than one earthworm replicate individual within the control (a, b; replicates Con 1, Con 2, Con 3) and cocktail (c, d; replicates Coc 1, Coc 2, Coc 3) treatments on nutrient agar (NA; a, c) and potato dextrose agar (PDA; b, d). The numbers in the brackets are the number of ASV representatives within each genera

## 4. Discussion

Earthworms are key soil organisms contributing to ecosystems processes and associated services [50]. It is recognised that earthworms harbour an abundant and diverse microbiome [51]. However, there are considerable uncertainties regarding the role of the earthworm microbiome in providing nutritional and non-nutritional benefits to the host and the consequences of the earthworm host-microbiome interaction for ecosystem processes such as OM decomposition. In this study we investigated the potential of antibiotics to create axenic earthworms for subsequent use in experiments aiming to improve our understanding of the role that the earthworm microbiome plays in host health and function. Previous studies have been carried out to produce axenic earthworms in *Eisenia fetida* [32,33]. However, the applicability of this method to species that can be considered more typical soil inhabitants was uncertain. Accordingly, here we extend the previous studies to consider species representing different earthworm ecotypes covering epi-anecic (*L. terrestris*) and epi-endo-anecic (‘intermediate’; *A. chlorotica*) ecological groups (according to Bottinelli et al.’s [6] re-classification).

As well as examining the impact of the various antibiotic treatments on the earthworm-associated microbial abundance, we additionally report on the diversity (richness) and composition of the culturable microbiome of *L. terrestris* and its response to antibiotic treatment.

Overall, we have shown that is it possible to significantly reduce the abundance of earthworm-associated culturable microorganisms through the treatment of earthworm individuals with antibiotics or antibiotic cocktail. Our approach is suitable for use in *E. fetida* and the soil dwelling species *L. terrestris* and *A. chlorotica*. However, the efficacy of antibiotic treatment depended upon the antibiotic(s) used, the earthworm species, and the agar medium used for microbial enumeration. In relation to the agar medium, we noted that colonies forming on PDA, like those for NA, were small and smooth resembling bacterial growth. Although PDA is associated with the cultivation of fungi (not bacteria), the composition of the medium (potato extract, glucose) does not select against bacterial growth. It contains glucose as a readily utilised C source. Given this colony appearance and also the observation that CFU abundance on PDA was not affected by the antifungal cycloheximide treatment (Figure 1), we assumed that colonies forming on PDA were bacterial.

Only the cocktail of five antibiotics (ampicillin, ciprofloxacin, cycloheximide, gentamicin and nalidixic acid) resulted in culturable numbers significantly lower than the control for both NA and PDA agar across all earthworm species (Figure 1), whilst ampicillin and cycloheximide mostly showed no significant differences when compared to the control. Resistance to ampicillin, a beta-lactam antibiotic, is known to be naturally prevalent among soil bacteria [52,53], and cycloheximide, an antifungal, is expected not to be effective on most bacteria [54].

It was possible, however, through the treatment of *E. fetida* (NA) and *A. chlorotica* with the antibiotic cocktail to reduce the abundance of earthworm-associated microorganisms from ≥ 10^5^ CFU per earthworm individual to below the limit of detection (50 CFU/ earthworm in our study). This agrees with previous studies [32,33] that have also applied antibiotics to create earthworms (*E. fetida*) deemed ‘axenic’ with no associated microorganisms detectable by culture.

Whilst the application of the antibiotic cocktail [and ciprofloxacin applied singly for *E. fetida* (PDA)] reduced culturable microbial abundance to below detection in *E. fetida* and *A. chlorotica*, microbial numbers were not reduced to below detection limits for *L. terrestris*, although a significant >100-fold knockdown was recorded in this species for the cocktail. To be exposed to antibiotics, through both dermal and gut contact, earthworm individuals needed to burrow and ingest agar. Differences in burrowing behaviour between species may influence the degree to which earthworms are exposed to antibiotics, and therefore the effectiveness of the antibiotic treatment. Also, there may also be different exposure levels in different bacterial populations. Bacteria in the gut are likely to receive a high dose, whereas the nephridial symbionts may be more ‘protected’ against antibiotics due to their embedment in an organ that may be more ‘sealed’ from antibiotics. *L. terrestris*’s natural behaviour is to create permanent vertical burrows, travelling to the surface to feed on partially decomposed plant litters and other organic matters [55,56]. Although we scaled up agar volumes to accommodate the larger *L. terrestris* size and burrowing behaviour, it is possible that *L. terrestris* individuals did not explore and ingest the antibiotic-containing agar to the same extent, resulting in reduced exposure. Alternatively, the *L. terrestris* microbiome may harbour a larger number of culturable antibiotic-resistant microorganisms [57,58]. Earthworms are known to produce their own antimicrobial agents [57] which might lead to earthworm species-specific selection of antibiotic resistance traits within the microbiome.

Although based on methods of Hand & Hayes[32] and Whiston and Seal [33], our approach differed from previously published work in terms of the spectrum of antibiotics applied. Nalidixic acid, gentamicin, a penicillin (ampicillin) and cycloheximide [33] or cycloheximide [32] were in common with the previous studies, but, additionally ciprofloxacin (a fluoroquinolone) was included as an antimicrobial not tested previously. In most cases ciprofloxacin, when applied alone, was just as effective in reducing culturable numbers as the cocktail treatment. This effectiveness may be related to its broad-spectrum DNA gyrase inhibitory activity against both Gram-negative and Gram-positive bacteria [59]. Nalidixic acid similarly inhibits bacterial DNA gyrase [60,61] whereas gentamicin has a different mode of action making it effective only towards Gram-negative bacteria [62].

As well as differences in the choice of antibiotics used, our method also varied from previously published work in terms of the methodology and duration of antibiotic exposure. We used sterile semi-solid technical agar as the ‘carrier’ for antibiotic exposure. In contrast, previous studies used aqueous solutions [32] or sterile suspensions of microcrystalline cellulose [33]. Our exposure period (4 days) was comparable to that employed by Whiston and Seal [33] (5 days), but shorter than the 35 days adopted by Hand and Hayes [32] and consisted of a single exposure step as opposed to one [33] or several [32] transfers of earthworm individuals between different antibiotic-containing media. Reducing the timescale of exposure and the degree of earthworm handling reduces the risk of earthworm mortality and ensures that an earthworm goes forwards in an unstressed state into further experiments. In our trial, all earthworm specimens survived after the exposure to the antibiotic when using response to touch stimuli as a superficial measurement of health condition.

For *L. terrestris*, 16S rDNA amplicon sequencing of the NA- and PDA-grown bacterial communities was applied to characterise the richness and composition of the culturable microbiome of *L. terrestris* and its response to antibiotic treatment. For reasons previously discussed, PDA-grown colonies were assumed to be bacterial and were included in the 16S rDNA-based sequencing effort. This enabled the characterisation of potentially different, agar specific, microbiomes due to the selective nature of bacterial growth [63].

Whilst cognisant that the bacteria that can be cultured on laboratory media are only a very small proportion of the total diversity and therefore that we have not captured what might be a significant non-culturable fraction [64], we focussed on culturable microbiomes (i.e., amplicon sequencing from colony-extracted DNA). This was because we were concerned that amplification of microbial DNA directly extracted from earthworm tissues would not be able to distinguish between DNA from living microbial cells surviving the biocidal treatments and those that had been recently killed [65]. Amplification of DNA from dead microorganisms would undermine the identification of bacterial taxa that escaped the effect of the antibiotic treatment. Since this culture-based approach will mean that the relative abundance of a given ASV in a sample will depend not only on the original cell abundance in the earthworm sample but also confounded by the subsequent rate of multiplication on agar, the subsequent analysis of diversity and taxonomic composition was based on presence/absence, not relative abundance.

Comparison of estimated Chao1 ASV richness and Observed richness suggested that the sequencing depth covered the richness of ASVs present. However, there was substantial within-treatment variation in ASV richness, including for the non-antibiotic-treated controls. Due to the destructive nature of sampling, it was not possible to examine the impact of antibiotic treatment on the microbiome for a given earthworm individual (i.e., before and after treatment). That there was no significant effect of antibiotic treatment on either ASV richness (Observed and Chao1) or phylogenetic richness, even for antibiotic treatments that significantly reduced the number of culturable bacteria (Cocktail (NA & PDA) and ciprofloxacin (NA); Figure 1), might be partly due to initial variability in bacterial richness between earthworm individuals (Figure 1) going into the incubation. This variability is in agreement with other reports of high variability in host -associated microbiomes [66–68]. When compared to other studies on earthworm-associated bacterial richness [67–69], our analysis revealed a low number of ASVs per worm (e.g., ~30 ASVs for the NA control). However, this is expected due to the focus on only those bacteria that formed colonies on the NA and PDA medium. In addition, the *L. terrestris* individuals in the current trial were depurated before the plating of earthworm samples. This means that the culturable microbiome in our study was likely not dominated by diverse transient microbes associated with the ingested loam: peat substrate but those more tightly-associated with the gut and other organs [68].

Whilst there was no significant impact on the richness of ASVs, PERMANOVA and ANOSIM analysis suggested an impact of antibiotic treatment on community composition. The PCoA (Figure 3) highlighted the variability between within-treatment replicates but suggested that the bacterial community compositions for the antibiotic treatments (cocktail, ciprofloxacin) that caused the most significant reduction in culturable abundance (Figure 1) were among the most dissimilar to the control.

Genera common to more than one earthworm individual within the same treatment were visualized by Venn diagrams (Figure 4) to identify core members of the culturable microbiome and those genera sensitive or tolerant to antibiotic treatment. The core bacterial diversity (phyla level) of the *L. terrestris* culturable microbiome composed of members of the *Proteobacteria* (*Aeromonas, Comomonas, Kosakonia, Lelliottia, Pseudomonas, Raoultella, Verminephrobacter spp*.) and *Bacteroidetes* (*Pedobacter, Sphingobacterium spp*.). This composition is in broad agreement with the earthworm-associated phyla described in other earthworm microbiome studies [70–72]. In particular, members of the genus *Verminephrobacter* are known symbionts found in Lumbricid earthworms and have a known nephridial association [31,73,74]. *Aeromonas*, a genus consisting of facultative anaerobic species, are a further taxa that are frequently earthworm-associated including with *L. terrestris* [75,76].

Among taxa indicating potential resistance, the near ubiquitous detection of *Aeromonas* in the culturable microbiome of both control and antibiotic cocktail treated individuals suggests that representatives of this genus were resistant to antibiotic treatment. *Aeromonas* are considered to be naturally resistant to *β*-lactam antibiotics, such as ampicillin [77,78] and resistance to ciprofloxacin and nalidixic acid has also been reported for environmental strains, including multi-antibiotic resistance [78]. In contrast, resistance of this genera to gentamicin appears to be rare [78,79]. Further characterization of the antibiotic resistance profile of our *Aeromonas* isolates would be required to discern if these strains were indeed gentamicin resistant as may be suggested by their presence in the cocktail exposure or, alternatively, evaded exposure. *Aeromonas hydrophila* has been isolated from the coelomic cavity of *L. terrestris* [80]. If *Aeromonas* were in this compartment, their exposure may be more limited than for bacteria in the gut. The organ-specific location of *Verminephrobacter* may similarly result in a lower exposure for members of this genus.

In contrast to the apparent tolerance of *Aeromonas* species to the antibiotic exposure, bacteria belonging to the genera *Comomonas, Kosakonia* and *Sphingobacterium* that were part of the core in control *L. terrestris* were not detected in cocktail-treated individuals. This absence suggests a possible antibiotic sensitivity of these strains. No antibiotic resistance genes have been annotated in environmental isolates of *Comamonas* [81] and we could not find reports of resistance traits in environmental *Kosakonia* and *Sphingobacterium* strains. The genus *Pseudochrobactrum*, however, was not detected in control individuals but was present in all cocktail-treated individuals (NA) suggesting that antibiotic treatment potentially promoted the growth of this putatively multi-antibiotic resistant group to densities above the limit of detection of the spread plate. We could not find any previous descriptions of the resistance of *Pseudochrobactrum* to the antibiotics used here. Further characterization is required to verify the antibiotic resistance profile of this group and to explore the earthworm as a microbial environment conducive to acquisition of antibiotic resistance genes, particularly under the pressure of antibiotic selection [82].

### 4.1 Conclusion

We have shown that is it possible, across three ecologically-contrasting earthworm species (*E. fetida, L. terrestris, A. chlorotica*), to significantly reduce the abundance of earthworm-associated culturable microorganisms through a 96 h exposure of earthworm individuals to a cocktail of antibiotics containing cycloheximide (150 μg ml^−1^), ampicillin (100 μg ml^−1^), ciprofloxacin (50 μg ml^−1^), nalidixic acid (50 μg ml^−1^), and gentamicin (50 μg ml^−1^)) in a semi-solid agar carrier. Abundance was reduced to below detection limits (50 CFU individual^−1^) in *E. fetida* and *A. chlorotica* and by >100-fold for *L. terrestris* with accompanying shifts in *L. terrestris* bacterial community composition. The culturable bacterial microbiome of control (non-exposed) and antibiotic cocktail-exposed *L. terrestris* individuals revealed between-individual variability in richness and diversity but also ‘core’ genera that were putatively sensitive (*Comomonas, Kosakonia* and *Sphingobacterium*) or resisted (*Aeromonas, Pseudochrobactrum*) antibiotic exposure. This characterization of the efficacy of antibiotic treatment in creating ‘axenic’ *E. fetida* and *A. chlorotica* individuals or *L. terrestris* with a supressed microbial composition provides the foundation for future experiments aimed at understanding the importance of earthworm-associated microorganisms, be they transient gut inhabitants or more permanently-associated, for host health and ecosystem functioning.

## 6. Acknowledgements

This study acknowledges PhD studentship funding from the Scenario NERC Doctoral Training Partnership grant NE/L002566/1.

